# Novel Interaction between Prefrontal and Parietal Cortex during Memory Guided Saccades

**DOI:** 10.1101/2020.03.11.985259

**Authors:** Nathan J. Hall, Carol L. Colby, Carl R. Olson

**Author notes:** Author contributions: All authors participated in design of the experiment, analysis of the data and writing of the paper. N.J.H. performed the research. Corresponding author: Carl Olson, Center for the Neural Basis of Cognition, 115 Mellon Institute, 4400 Fifth Avenue, Pittsburgh, PA 15213. NJH: Department of Neurobiology, Duke University, Box 3209, Durham, NC 27710.

## Abstract

Dorsolateral prefrontal cortex (DLPFC) and posterior parietal cortex (PPC) are linked to each other by direct reciprocal connections and by numerous pathways that traverse other areas. The nature of the functional coordination mediated by the interconnecting pathways is not well understood. To cast light on this issue, we simultaneously monitored neuronal activity in DLPFC (areas FEF and 8a) and PPC (areas LIP and 7a) while monkeys performed a memory guided saccade task. On measuring the spike-count correlation, a measure of the tendency for firing rates to covary across trials, we found that the DLPFC-PPC correlation became negative at the time of the saccade if and only if the neurons had matching spatial preferences and the target was at their mutually preferred location. The push-pull coordination underlying the negative spike-count correlation may help to ensure that saccadic commands emanating from DLPFC and PPC sum a constant value.

**Significance:** Anatomical pathways linking cortical areas that mediate executive control are thought to mediate coordination between them. We know very little, however, about the principles that govern this coordination. In the present study, we addressed this issue by recording simultaneously from neuronal populations in prefrontal and parietal cortex while monkeys performed memory guided saccades. We found a clear sign of coordination. Prefrontal and parietal neurons encoding a given saccade engage in a push-pull interaction during its execution. If parietal neurons are more active, prefrontal neurons are less active and vice versa. We suggest that this is a manifestation of a general principle whereby commands emanating from DLPFC and PPC are coordinated so as to sum a constant value.

## Introduction

Dorsolateral prefrontal cortex (DLPFC) and posterior parietal cortex (PPC) are interconnected by strong topographically organized reciprocal pathways (Cavada and Goldman-Rakic, 1989; Andersen et al., 1990; Schall et al., 1995; Stanton et al., 1995; Rozzi et al., 2006) and share connections to a common set of other areas (Selemon and Goldman-Rakic, 1988). The numerous connections between them presumably mediate some form of coordination in the performance of functions dependent on their combined activity. One context in which coordination presumably occurs is the memory guided saccade (MGS) task (Hikosaka et al., 1989). Neurons active during MGS performance occupy a swath of DLPFC encompassing the frontal eye field (FEF) and anteriorly adjacent cortex (area 8a) and a territory in PPC encompassing the lateral intraparietal area (LIP) and posteriorly adjacent cortex (area 7a). DLPFC and PPC neurons exhibit nearly identical patterns of activity during visual, delay-period and saccadic epochs of the MGS task (Funahashi et al., 1989; Barash et al., 1991; Colby et al., 1996; Chafee and Goldman-Rakic, 1998; Katsuki and Constantinidis, 2012a) and in other tasks as well (Buschman and Miller, 2007; Qi et al., 2010; Merchant et al., 2011; Goodwin et al., 2012; Katsuki and Constantinidis, 2012b; Zhou et al., 2012; Suzuki and Gottlieb, 2013; Katsuki et al., 2014b; Katsuki et al., 2014a; Qi and Constantinidis, 2015; Sarma et al., 2016; Zhou et al., 2016). However, the contributions of DLPFC and PPC to MGS performance are not identical. DLPFC is in stronger and more direct control of saccadic output than PPC as evidenced by the observations that inactivation of DLPFC produces a more severe saccadic impairment than inactivation of PPC (Dias and Segraves, 1999; Li et al., 1999), that electrical stimulation of DLPFC produces saccades at lower current threshold than electrical stimulation of PPC (Shibutani et al., 1984; Bruce et al., 1985; Kurylo and Skavenski, 1991; Thier and Andersen, 1996), and that DLPFC, unlike PPC, sends direct projections to pre-oculomotor pontine nuclei (Leichnetz et al., 1984a; Leichnetz et al., 1984b).

The most straightforward way in which to characterize coordination between DLPFC and PPC is to determine how neuronal activity in one area depends on neuronal activity in the other. Intervention-based studies have provided evidence for dependency. Electrical stimulation of DLPFC affects saccade-related activity in PPC (Premereur et al., 2012; Premereur et al., 2014) and cooling of each area affects saccade-related activity in the other (Chafee and Goldman-Rakic, 2000). Correlation-based studies have provided further evidence for dependency. Phase-locking of local field potential oscillations has been observed in the context of a visual search task (Buschman and Miller, 2007) and an object working memory task (Salazar et al., 2012; Dotson et al., 2014). Likewise, in an analysis of the resting state BOLD signal, synchrony has been observed on long time scales (Hutchison et al., 2012). While these studies have established the interdependence of neural processes in DLPFC and PPC, they have left open an important question regarding the computational significance of the interactions: is the pattern of dependence related to the functional properties of the interacting neurons? A single previous study, concerned with population dynamics, has yielded evidence of interactions dependent on neuronal spatial selectivity in a spatial categorization task (Crowe et al., 2013).

The MGS task provides an ideal context in which to characterize DLPFC-PPC interactions dependent on neuronal spatial selectivity because neurons in both regions are selective for saccade direction. Accordingly, we set out to measure coordination between DLPFC and PPC at the level of single neurons in the MGS task. We analyzed cross-neuronal coordination by use of a measure, the spike-count correlation or r_sc_, sensitive to the tendency for the activity of two neurons to covary across identical trials. This approach has often been applied to neurons in the same area (Cohen and Kohn, 2011) but it has been applied less frequently to neurons in different areas (Pooresmaeili et al., 2014; Oemisch et al., 2015; Ruff and Cohen, 2016a, b).

## Materials and Methods

### Subjects

Two adult male rhesus monkeys (*macaca mulatta*) were used in these experiments. They were cared for in accordance with National Institutes of Health guidelines. The Institutional Animal Care and Use Committees of Carnegie Mellon University and the University of Pittsburgh approved all experimental protocols. Monkeys CY and RY weighed 13.0 and 8.0 kg respectively.

### Experimental apparatus

The monkey sat in a primate chair with head fixed in a darkened room viewing a CRT monitor at a distance of 30 cm (19” ViewSonic^**®**^ color CRT monitor at a refresh rate of 85 Hz using an 8 bit DAC with an ATI Radeon™ X600 SE graphics card). Stimulus presentation, monitoring of eye position and delivery of reward were under the control of NIMH Cortex software (provided by Dr. Robert Desimone). Eye position was monitored with an infrared eye tracker sampling at 240 Hz (ISCAN Inc., Woburn, MA). Eye position voltage signals were continuously monitored and saved at a sampling rate of 1000 Hz for offline analysis on a separate computer running Plexon software (Plexon Inc., Dalls, TX). Data analysis was carried out offline using custom MATLAB^**®**^ software (Mathworks, Natick, MA).

### Chamber placement

Each monkey was equipped with a surgically implanted plastic cranial cap that held a post for head restraint and two cylindrical recording chambers 2 cm in diameter. These were oriented normal to the cortical surface with the base of the frontal chamber centered over the genu of the arcuate sulcus and the base of the parietal chamber centered over the intraparietal sulcus. The chambers were positioned over the left hemisphere in monkey CY and the right hemisphere in monkey RY. Their placement was guided by MR images showing gray matter and white matter together with fiducial markers placed at known locations within the cranial implant. Electrodes were advanced into the cortex underlying each chamber along tracks forming a square grid with 1 mm spacing.

### Memory guided saccade task

The monkeys were trained to perform a memory guided saccade (MGS) task. At the beginning of each trial, the monkey maintained fixation on a 1°x1° fixation cross for a randomly selected interval in the range 300-500 ms. Then, as the monkey continued to fixate, a white circle 0.5° in diameter appeared in the visual field periphery for 47 ms (four video frames). The monkey continued to maintain central fixation during an ensuing delay period with a randomly selected duration in the range 400-1200 ms. At the end of this interval, offset of the fixation cross signaled the monkey to make a saccade to the remembered location of the target. The monkey was required to execute a saccade into a 3°x3° window centered on the target, exiting the central window within 500 ms and entering the target window within an additional 120 ms. The target reappeared when the gaze entered the window. The monkey was required to maintain gaze within the window for an additional period randomly selected from the range 200-400 ms. Successful completion culminated in delivery of liquid reward. On interleaved trials, the target could appear at multiple locations. The number of locations varied according to the phase of the experiment (preliminary mapping, selection of data-collection sites, data collection) as described below. The sequence of locations across trials was random subject to the constraint that the target be presented with equal frequency at each location.

### Preliminary mapping

Before the phase of the experiment involving data collection, we mapped out functionally defined areas underlying the two chambers using single tungsten microelectrodes (Frederick Haer Company). We measured neuronal activity in the context of the memory guided saccade task, presenting targets at twelve locations spaced evenly around the clock at various eccentricities. In frontal cortex, we assessed whether intracortical microstimulation elicited eye movements. In parietal cortex, we checked whether neurons were active in conjunction with limb movements or responded to manually delivered visual and somatosensory stimuli. The frontal eye field (FEF) was defined as consisting of sites within the anterior bank of the arcuate sulcus and on the adjacent gyrus at which neurons exhibited spatially selective visual and saccadic responses in the context of the memory guided saccade task and at which saccades could be elicited with electrical stimulation (bipolar train, 250 µs pulse width, 350 Hz, 100 ms duration) at currents less than 50 µA (Bruce et al., 1985). The lateral intraparietal area (LIP) was defined as consisting of sites within the lateral bank of the intraparietal sulcus, adjacent to the ventral intraparietal and medial intraparietal areas (Colby et al., 1996), at which neurons exhibited spatially selective visual and saccadic responses during performance of the memory guided saccade task. The lateral boundary of LIP was indeterminate on functional grounds as sites in area 7a close to the lip of the intraparietal sulcus and on the adjacent gyrus also exhibit spatially selective task-related activity.

### Data collection

We monitored neuronal spiking activity through two 8-channel linear microelectrode arrays, one in DLPFC and one in PPC, with recording sites distributed along the shaft at intervals of 150 μm (Alpha Omega Co. USA Inc., Alpharetta, GA). Occasionally, we substituted for the linear array in PPC a tungsten microelectrode with a single contact (Frederick Haer, Bowdoinham, ME). At the beginning of each day’s session, the electrodes were introduced simultaneously into the frontal and parietal cortices through stainless steel guide tubes stabilized in a nylon grid system (Crist Instrument Co. Inc., Hagerstown MD). As we advanced the electrodes, we monitored neuronal activity while the monkey performed a version of the memory guided saccade task in which targets were presented at twelve locations on interleaved trials. The locations were of equal eccentricity and were distributed around the clock at 30° intervals.

Assessing responses at fixed eccentricity allowed characterizing preferred directions. Adjusting eccentricity between blocks allowed characterizing preferred amplitudes. We proceeded to data collection only if neurons on some channels exhibited spatially selectivity visual or perisaccadic activity. Neural activity on each channel was thresholded at 2-3 standard deviations above mean background noise. Threshold-crossing events alone were stored. These were amplified, filtered, and saved at a sampling rate of 40 kHz using Plexon MAP system hardware and software. Spike waveforms were sorted online and offline using Plexon software. We distinguished single-neuron waveforms from small-amplitude multi-unit activity (MUA) on the basis of their forming well-defined clusters in principal component space.

### Database

Before proceeding to data analysis, we winnowed the data set down to cases suitable for further analysis. This involved eliminating neurons with unsuitable functional properties and eliminating trials in which behavior or neuronal activity was aberrant. *1) Initial data set*. Out of 102 sessions, 7 yielded neural data from DLPFC alone, 7 yielded neural data from PPC alone and 88 yielded neural data from both areas. In DLPFC, the number of channels carrying MUA during a successful session ranged from 1 to 8 with a mean of 4.0 while the number of well-isolated spikes ranged from 0-3 per channel with a mean of 0.5. In PPC, the number of channels carrying MUA during a successful session ranged from 1 to 8 with a mean of 3.3 while the number of well-isolated spikes ranged from 0-3 per channel, with a mean of 0.6. In analyzing the data, we treated the MUA on each channel as if it emanated from a single neuron. We accordingly apply the term “neuron” to both MUA and well-isolated spikes.

Restricting consideration to well-isolated spikes did not affect the outcome of the experiment. *2) Elimination of neurons lacking perisaccadic activity*. We required, as a basis for including a neuron in the database, that its perisaccadic firing rate be greater, under at least one of the two target conditions, than its firing rate during a 300 ms baseline epoch immediately preceding target onset (Wilcoxon rank sum test, one-sided p < 0.10). The analysis window was aligned to the time of saccade onset defined as the moment at which the velocity of the eye first exceeded 30°/s. The window was centered on the period associated with maximal population activity in the area in question. For DLPFC, it extended from 100 ms before to 50 ms after saccade onset. For PPC, it extended from 50 ms before to 100 ms after saccade onset. *3) Elimination of trials involving aberrant saccades*. Trials were eliminated from the database if the behavioral reaction time, the saccade vector angle, or the saccade vector amplitude was more than 2 standard deviations from the session mean for trials with the target at the location in question. Elimination of a trial meant elimination of data collected from all neurons during that trial. *4) Elimination of trials involving low or deviant perisaccadic firing rates*. In the event that a neuron’s perisaccadic firing rate underwent a step change during the session, we eliminated from that neuron’s database the entire suspect block of trials. In the case of a decrease, indicating loss of the neuron, trials following the step were removed. In the case of an increase, indicating acquisition of the neuron, trials preceding the step were removed. If the neuron’s average firing rate was less than 1 Hz over a period of several minutes, then trials during the low-firing-rate period were removed. If a neuron’s perisaccadic firing rate on a given trial was more than 3 standard deviations from the neuron’s mean firing rate on trials with the target at the location in question, then the trial was removed from that neuron’s database. *5) Elimination of neurons with too few trials per condition*. If, after the steps described above, any neuron had fewer than 20 trials remaining for either of the two target locations, then that neuron was removed from the database.

### Spike-count correlation analysis

In every spike-count correlation analysis, we computed r_sc_, as the Pearson’s correlation coefficient between the firing rates of the two neurons across trials in which the target was at the same location. The duration of the window in which firing rate was computed was always 200 msec. Before each analysis, we reduced the database and conditioned the firing rates to remove the influence of extraneous covariates. We based these steps on neuronal activity in the window within which the spike-count correlation was to be computed. *1) Reducing the database*. We winnowed from the database any neuron pair for which, within the selected analysis window under the selected target condition, the average raw firing rate of either member was < 2 Hz or the geometric mean of the two raw firing rates was < 5 Hz. If the analysis concerned pairs of neurons recorded on the same linear array, then, upon completion of the analysis, if r_sc_ > 0.8, we excluded the pair from consideration on the ground that this might be a case in which the same spike was recorded on two channels. *2) Removing the influence of extraneous covariates*. Behavior and its context varied subtly from trial to trial even when the location of the target was the same. Neurons in DLPFC and PPC consequently could have exhibited a significant spike-count correlation because their firing rates were jointly locked to some extraneous covariate. To minimize the likelihood of such an effect, before carrying out each spike-count correlation analysis, we conditioned the raw firing rate of each neuron by the following procedure. We first square-root transformed the spike count so as to stabilize the variance (Yu et al., 2009). Then we regressed the square-root-transformed spike count on the following factors that varied across trials. *Time*: the sequential number of the trial within the run. *Delay*: the length of the preceding delay period. *Theta and Rho*: polar coordinates of the saccadic vector. Spike count correlation analysis was based on the firing-rate residuals remaining after removal of variance explained by these factors. *Start X and Start Y*: pre-saccadic gaze direction in Cartesian coordinates. *End X and End Y*: post-saccadic gaze direction in Cartesian coordinates. *Velocity*: peak velocity of the eye. *RT*: interval between offset of central fixation spot and initiation of saccade.

### Mean matching

Measurements of spike-count correlation can be affected by the firing rates of the neurons (de la Rocha et al., 2007; Cohen and Kohn, 2011). Consequently, if trial conditions differ with regard to the measured spike-count correlation, this could be an artifact of their differing with regard to firing rate. In the present study, this comment applies to the comparison between trials in which the saccade was directed to the location preferred by the neurons (yielding a high firing rate) and trials in which it was directed to the non-preferred location (yielding a low firing rate). The standard solution to this problem is to ask whether the difference between conditions with regard to spike-count correlation persists when comparison is confined to a subset of cases in which the geometric mean firing rate is equated across conditions (Cohen and Kohn, 2011). We implemented mean-matching in the following way. For each pair of neurons, independently for each condition, we measured both the spike-count correlation (r_sc_) and the geometric mean raw firing rate. Then we categorized the geometric mean firing rate observations into 1 Hz wide bins for each condition (Figure 5, *A*). The geometric-mean-firing rate distributions for the target-in and target-out conditions partially overlapped. If at least one observation from each condition fell into a bin, *i*, then we categorized that bin as belonging to the zone of overlap. This condition can be expressed as:

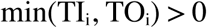

where TI_i_ was the number of observations in bin *i* for the target-in condition and TO_i_ was the number of observations in bin *i* for the target-out condition. Observations from all bins satisfying equation 1 were employed for resampling. On each of 10,000 iterations of the resampling procedure, we selected n pairs randomly with replacement from the pooled observations, where

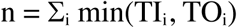

**Figure 1.**
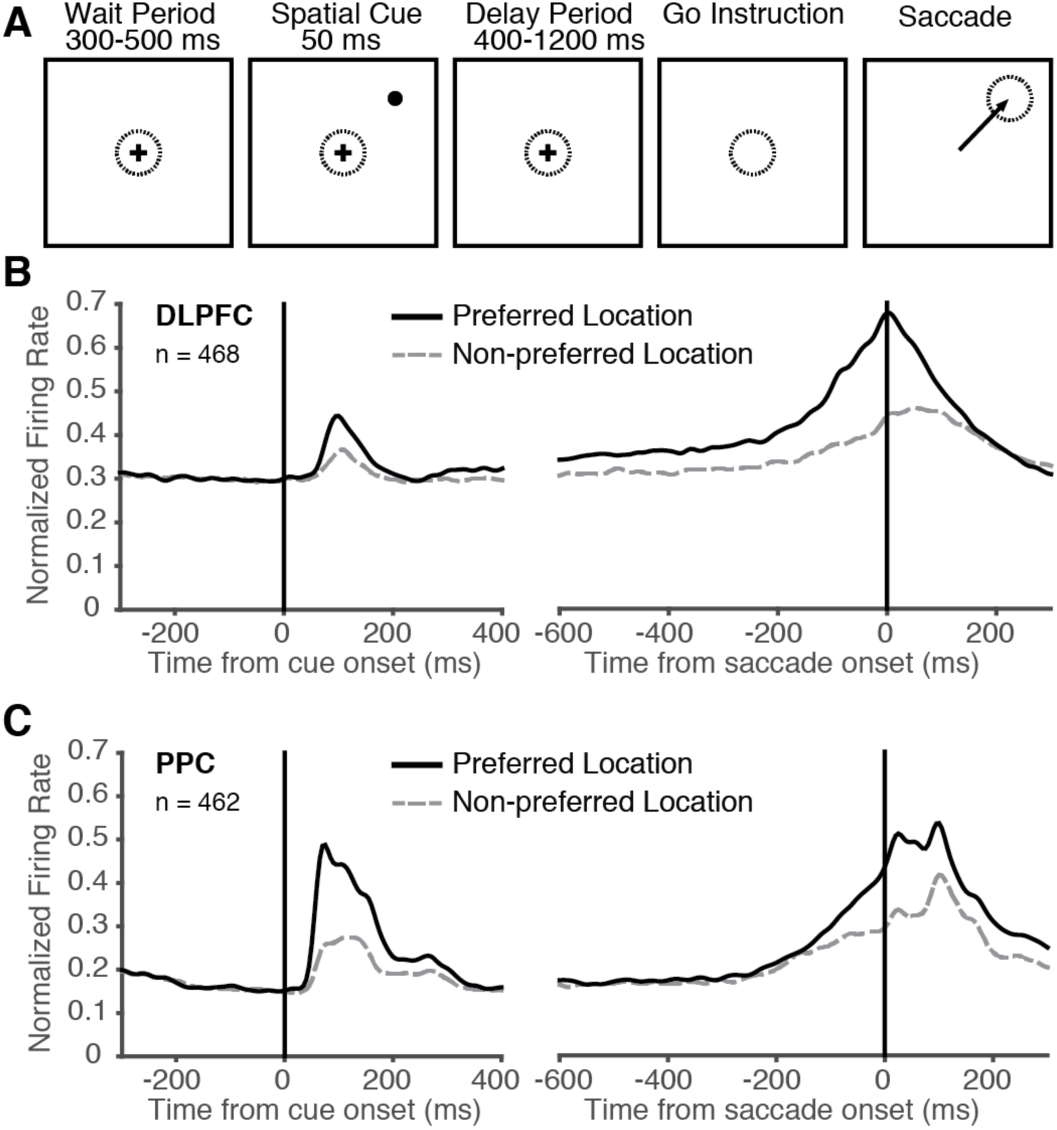
Population activity was spatially selective. ***A***, Sequence of events in a typical trial. Dashed circle is centered at the fovea. Arrow indicates saccade. The target could appear at either of two locations, one in the upper quadrant of the contralateral visual field and the other in the lower quadrant. ***B***, Mean population firing rate as a function of time during the trial for all neurons in the DLPFC database. Data sorted according to whether the target was at the preferred location (solid curve) or non-preferred location (dashed curve). The firing rate of each neuron was peak-normalized before the population average was computed. ***C***, Mean population firing rate as a function of time during the trial for all neurons in the PPC database.

**Figure 2.**
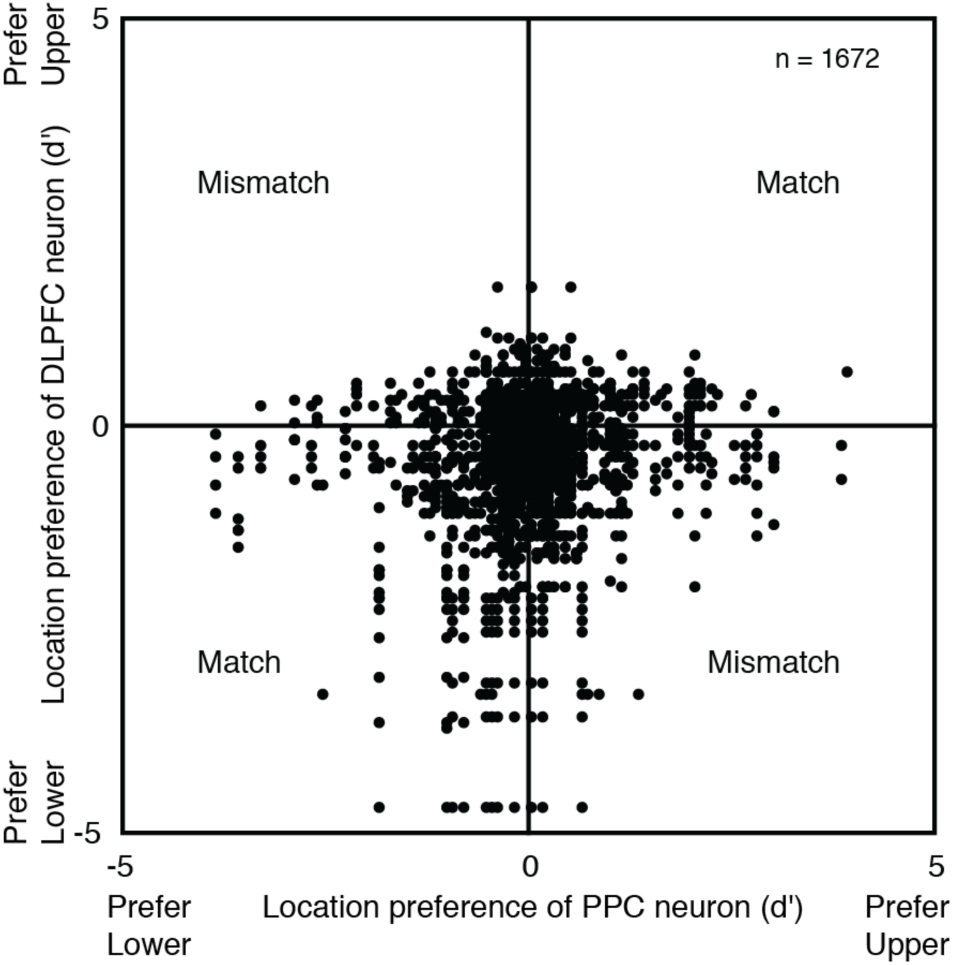
Neurons varied with respect to the degree of the preference for one location over the other. For each simultaneously recorded pair of neurons, the location preference index of the DLPFC neuron is plotted against the location preference index of the PPC neuron. Each measure is based on perisaccadic sensitivity to target location (d’) with d’ positive for the upper location and negative for the lower location. Note that low or high selectivity is not an absolute property of the neuron but rather a reflection of how poorly or well it differentiated between the two targets selected for use during the session. Cases in which the same neuron participated in multiple pairs account for the arrangement of points into rows and columns. Sixteen pairs containing at least one neuron with |d’| > 5 are excluded from the plot.

**Figure 3.**
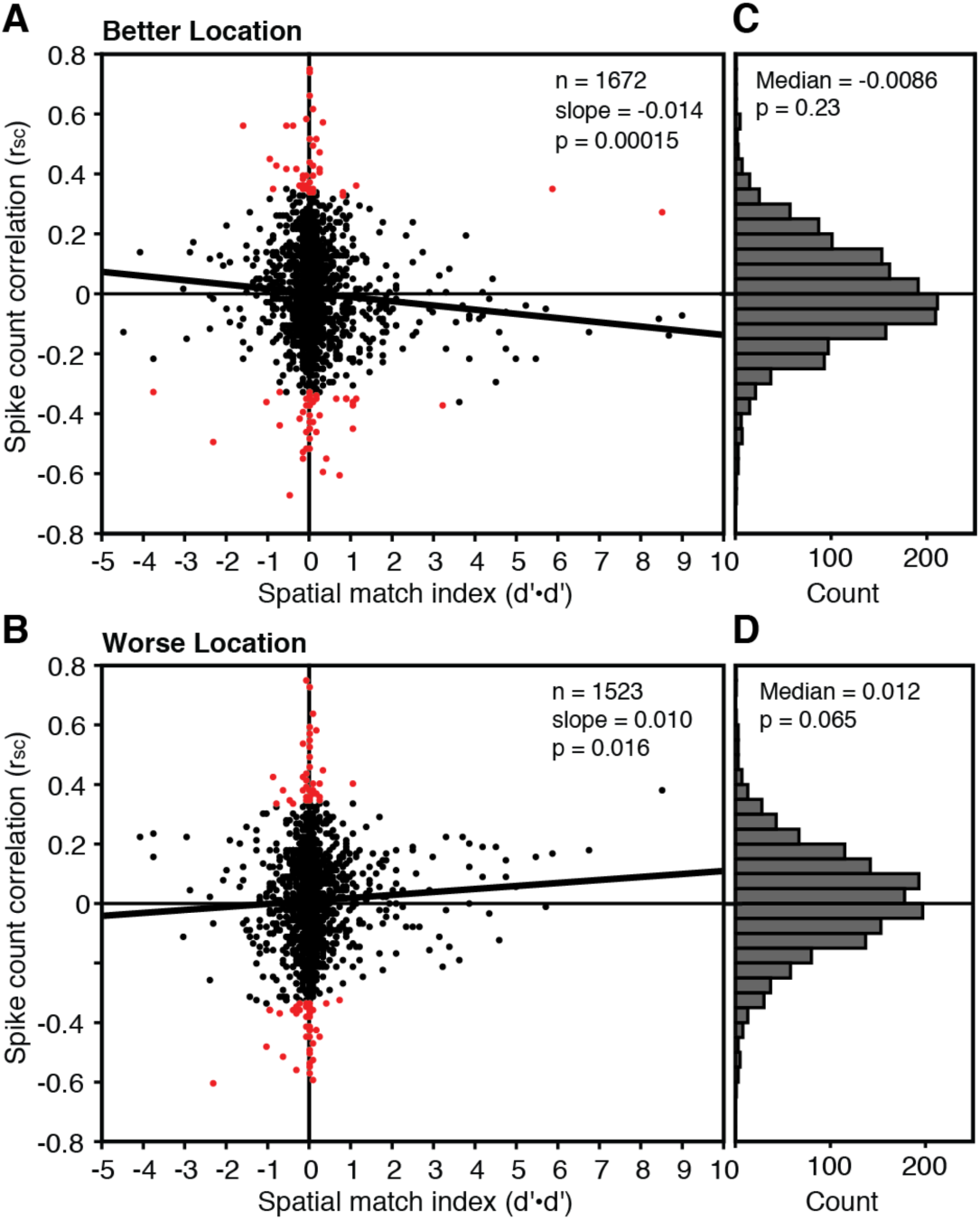
The perisaccadic spike-count correlation depended on the location preferences of the neurons and the location of the target. Each point represents a single DLPFC-PPC neuron pair. The perisaccadic spike-count correlation (r_sc_) is plotted against the spatial-match index (d’•d’) independently for cases in which the target was at the pair’s better, ***A***, or worse, ***B***, location as indicated by a higher or lower geometric mean perisaccadic firing rate. Red points indicate outliers discarded from the regression analysis. ***C-D***, Marginal distributions of spike-count correlations. The number of pairs satisfying the inclusion criteria for spike-count correlation analysis was greater for the better location (1672) than for the worse location (1523) because the firing rate was higher for the better location.

**Figure 4.**
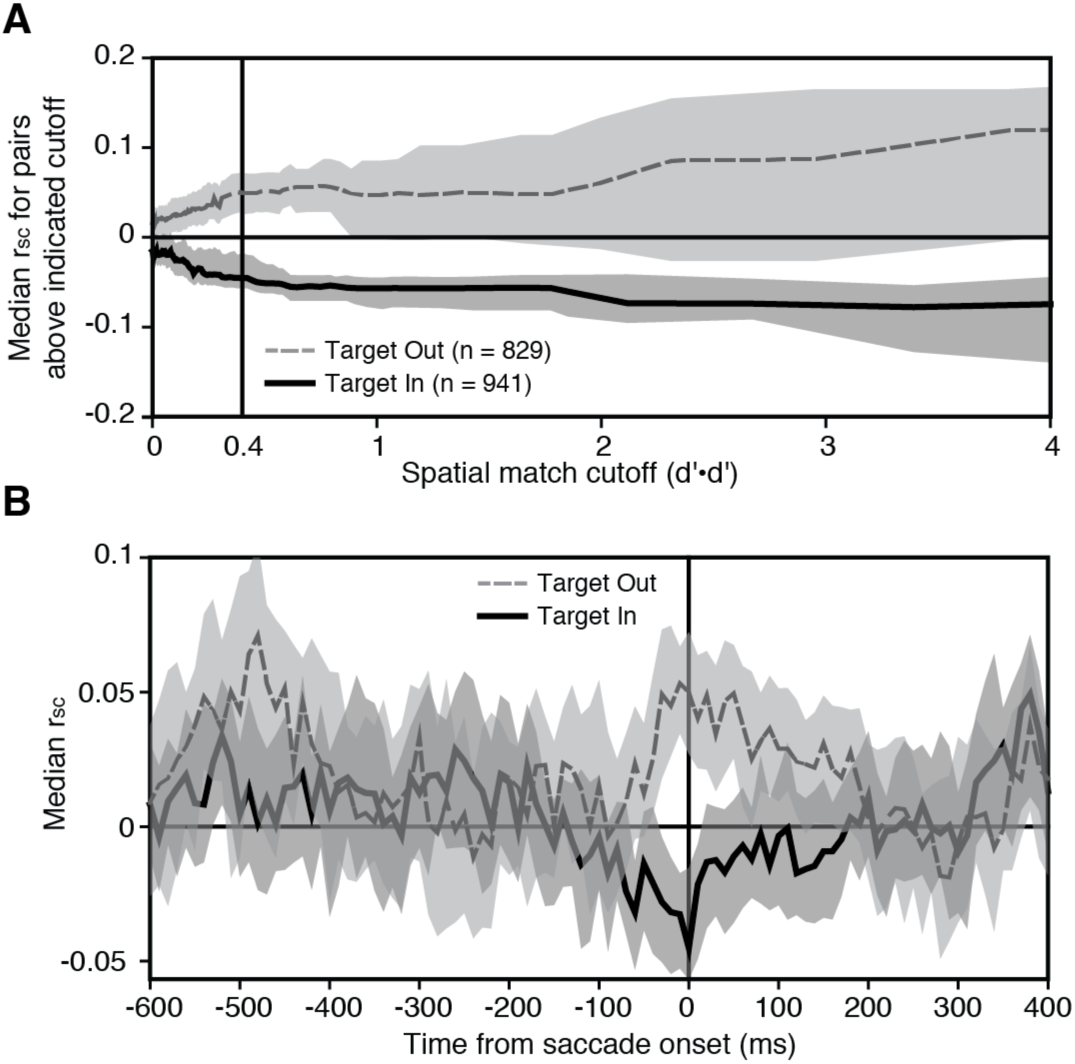
Neuron pairs strongly preferring the same location exhibited a push-pull effect during saccades to that location. ***A***, Median perisaccadic r_sc_ became negative on target-in trials as the subsample of neuron pairs was progressively narrowed (left to right) to exclude cases with low spatial-match index. Each curve connects points generated in the following manner. The population of neuron pairs was divided into percentiles on the basis of the spatial match index. Analyses were then carried out on the entire population of neuron pairs, on the subsample containing the top 99 percentiles, on the subsample containing the top 98 percentiles, and so on. The results of each analysis were represented by plotting the median r_sc_ of the subsample (vertical axis) against the spatial match index of the neuron pair with lowest rank in the subsample (horizontal axis). The curves are truncated for purposes of display at the right edge of the panel, excluding points for the best 2% and 1% of the population (target-in condition) and for the best 1% of the population (target-out condition). ***B***, Spike-count correlation as a function of time relative to saccade initiation. The analysis was based on firing in a 200 ms window with its center stepped in 10 ms increments relative to saccade initiation. It was restricted to neuron pairs with a spatial match index ≥ 0.4. These pairs numbered 293 for the target-in condition and 234 for the target-out condition. The sample in each window was further restricted to neuron pairs meeting, in that window, the criteria for inclusion in r_sc_ analysis. Ribbons in ***A-B*** indicate 95% bootstrap confidence intervals.

**Figure 5.**
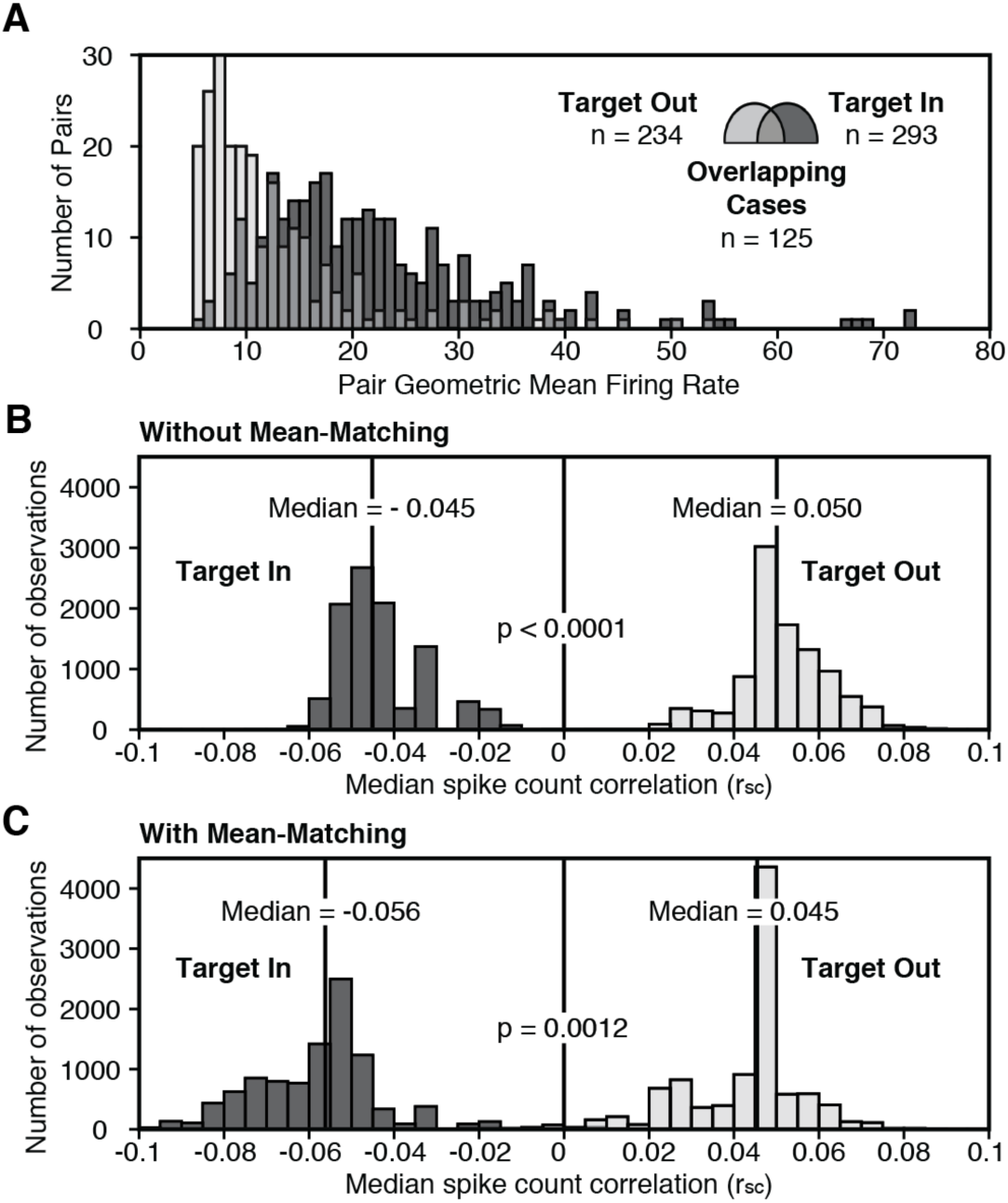
The push-pull effect survived factoring out the effect of mean firing rate. ***A***, The horizontal axis represents neuron-pair geometric-mean firing rate in a 200 ms window centered on saccade initiation. The frequency distribution of DLPFC-PPC neuron pairs under the target-in condition (dark bars) is displaced to the right relative to the frequency distribution of DLPFC-PPC neuron pairs under the target-out condition (pale bars). Each n indicates the count of neuron pairs with spatial match index ≥ 0.4 that met the conditions for r_sc_ analysis in a 200 ms perisaccadic window. The target-in and target-out distributions overlap in a zone of intermediate gray containing a summed count of 125. ***B***, Distribution of the median spike-count correlations (r_sc_) obtained by resampling from the full population of neuron pairs without regard to geometric mean firing rate. ***C***, Distribution of the median spike-count correlations (r_sc_) obtained by resampling pairs of observations, one from the target-in condition and one from the target-out condition, with matched geometric mean firing rates.

For each of the n randomly selected pairs, we selected a second pair randomly with replacement under two constraints: (1) the second pair was from the opposite condition and (2) the geometric mean firing rate of the second pair was from the same bin, *i*, as for the first pair. This procedure yielded two distributions of n neuron pairs, one for the target-in condition and the other for the target-out condition, that were matched with regard to geometric mean firing rate. We computed the median of r_sc_ for the target-in distribution and for the target-out distribution. Repeating this procedure 10,000 times yielded a distribution of 10,000 median r_sc_ values for the target-in condition and likewise for the target-out condition. As a basis for comparison, we carried out a parallel analysis in which we resampled observations from the entire target-in data set and likewise from the entire target-out data set without regard to geometric mean firing rate.

### Spatial selectivity index

To characterize each neuron’s pattern of spatial selectivity during the perisaccadic epoch, we computed d’ according to the following formula:

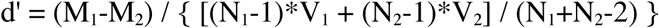

where M and V were the mean and variance of firing rate, and N was the number of trials. The subscripts 1 and 2 denote conditions in which the target was placed in the upper and lower quadrants respectively.

### Experimental design and statistical analysis

All statistical analyses were carried out in Matlab (https://www.mathworks.com/). Individual analyses are described in Results. The statistical tests used in these analyses, including bootstrap tests, the Wilcoxon signed rank test and linear regression with a large sample size, do not assume normality in the data.

## Results

We simultaneously monitored the activity of neurons in DLPFC and PPC of the same hemisphere while monkeys performed a memory guided saccade task (Figure 1, *A*). On randomly interleaved trials, the target appeared at one of two locations contralateral to the recording hemisphere. The locations were selected at the outset of the session to ensure that as many recorded neurons as possible exhibited spatially selective activity at the time of the saccade. One location was always in the upper quadrant and the other always in the lower quadrant and the two locations always subtended at least 90° at the fovea. In the average session, the monkey successfully completed 80 trials (range, 30-100) with the target at each location.

During most sessions, we monitored neuronal activity in each region through a linear microelectrode array containing eight contacts at 150 □m spacing; however, in a few sessions, the PPC electrode contained only a single contact. DLPFC recording sites were at grid coordinates coincident with FEF as identified during preliminary mapping and extending up to 3 mm anterior to it. PPC recording sites were at grid coordinates and depths coincident with LIP and laterally adjacent area 7a as identified during preliminary mapping. There were no obvious regional trends in either DLPFC or PPC with regard to the functional properties of neurons. Accordingly, we have not subdivided the data according to precise recording location.

We collected data during 109 sessions in two monkeys (66 in monkey CY and 43 in monkey RY). We selected for analysis all neurons that fired significantly more strongly during saccades to one target or both than during a pre-target baseline period. The 200 ms peri-saccadic window was centered 50 ms before saccade onset for DLPFC and 50 ms after saccade onset for PPC so as to center it at the time of maximal population response strength. This selection procedure yielded a total of 468 DLPFC neurons and 462 PPC neurons. As a population, these neurons carried time-varying spatially selective signals (Figure 1, *B-C*) consistent with those described in previous reports. The ensuing spike-count correlation analysis was based on 1672 simultaneously recorded DLPFC-PPC neuron pairs involving 428 DLPFC neurons and 395 PPC neurons.

We measured the perisaccadic spike-count correlation between neurons in each DLPFC-PPC pair for each saccade direction. The analysis was based on firing in a 200 ms window centered at saccade onset and was limited to data from trials, neurons and neuron pairs that met strict inclusion criteria and from which the influence of extraneous behavioral and contextual covariates had been factored out. To establish the statistical significance of the correlation for individual neuron pairs was not feasible because the number of trials was too small. Accordingly, statistical analysis focused on testing whether the median of the distribution across all neuron pairs was significantly different from zero.

At the coarsest level of pooling, which is to say in data combined across all neuron pairs under both target conditions, no trend was apparent. The median of the distribution of spike-count correlations was statistically indistinguishable from zero (median = 0.0021, p = 0.70, n = 3195, sign test). The absence of an effect might have arisen from combining data across cases in which the trends were of opposite sign. To investigate this possibility, we explored the dependence of the spike-count correlation on the preferred locations of the paired neurons and the location of the target. We characterized the spatial sensitivity of each neuron with a signal-detection-based measure (d’) which, by convention, was positive for upper-target preference and negative for lower-target preference. We took the multiple of the two d’ values as an index of the degree of match (if positive) or mismatch (if negative) between the spatial preferences of the paired neurons (Figure 2). This is a standard approach under circumstances in which the use of only a few locations prevents measuring signal correlation (Ruff and Cohen, 2014b). We characterized target location as better for the pair (more effective at eliciting perisaccadic firing) or worse for the pair (less effective at eliciting perisaccadic firing) on the basis of which produced the higher geometric mean firing rate. We regressed the spike-count correlation on the spatial match index independently for cases in which the target was at the neuron pair’s better or worse location. With the target at the better location (Figure 3, *A*), r_sc_ exhibited a significant negative dependence on the spatial match index (beta = - 0.014, p = 0.00015, n = 1672). This effect was driven by neuron pairs with a positive spatial match index (beta = -0.017, p = 0.00022, n = 941) and not by neuron pairs with a negative spatial match index (beta = -0.0016, p = 0.86, n = 731). With the target at the worse location (Figure 3, *C*), the spike-count correlation exhibited a significant positive dependence on the spatial match index (beta = 0.010, p = 0.016, n = 1523). This effect was driven by neuron pairs with a positive spatial match index (beta = 0.014, p = 0.0072, n = 829) and not by neuron pairs with a negative spatial match index (beta = -0.013, p = 0.22, n = 694).

We proceeded to ask whether, for neuron pairs with a large positive spatial match index, the spike-count correlation was significantly negative on target-in trials and significantly positive on target-out trials. To answer this question, we computed median r_sc_ repeatedly while progressively narrowing the pool of neuron pairs, first removing pairs with a spatial match index in the lowest percentile, then removing pairs with a spatial match index in the lowest two percentiles, and so on. As the pool narrowed, the refined subsample of neuron pairs came to exhibit a significantly negative spike-count correlation when the target was at the jointly preferred location and a significantly positive spike-count correlation when it was not (Figure 4, *A*). This pattern was well established for neuron pairs with a spatial match index ≥ 0.4 (vertical line in Figure 4, *A*). Beyond this level, both effects held steady although the confidence limits grew broader due to lessening of statistical power attendant on the reduction of the number of pairs. The negative spike-count correlation can be understood as arising from a push-pull effect whereby, if one neuron becomes more active the other becomes less active.

The analyses described up to this point were based on firing during the perisaccadic epoch. We next analyzed whether the results were specific to this epoch. To address this issue, we considered all neuron pairs with a spatial match index ≥ 0.4. For these pairs, we computed the median spike-count correlation in a 200 ms window with its center stepped in 10 ms increments from 600 ms before to 400 ms after initiation of the saccade. At around the time of saccade onset, the spike-count correlation underwent a negative excursion on target-in trials and a positive excursion on target-out trials (Figure 4, *B*). It is noteworthy that the spike-count correlation trended positive throughout the antecedent delay period under both conditions because it indicates that the pattern of interaction driving it into the negative range during target-in trials was specific to the time of saccade execution.

Differences in firing rate can affect the magnitude of the measured spike-count correlation (de la Rocha et al., 2007). The firing-rate on target-in trials was higher by definition than on target-out trials. Consequently, it was necessary to examine whether the difference between the target-in and target-out conditions would withstand removing the influence of firing rate. To resolve this issue, we carried out an analysis based on all neuron pairs with a spatial match index ≥ 0.4. For each pair, we computed the geometric mean of the perisaccadic firing rates. The distribution of geometric means was shifted to the right for the target-in condition as compared to the target-out condition, by definition, but there was a zone of overlap (Figure 5, *A*). Upon randomly resampling cases from the full distributions without any constraint on geometric mean firing rate, we found, as expected, that the median of the resampled perisaccadic spike-count correlations was negative under the target-in condition and positive under the target-out condition (Figure 5, *B*). We then repeated the resampling procedure, considering cases only from the zone of overlap and requiring that for each target-in case there be a target-out case with the same geometric mean firing rate. This mean-matching procedure yielded results virtually identical to those obtained without mean-matching (Figure 5, *C*). We conclude that the difference between target-in and target-out conditions with respect to the sign of the spike-count correlation was not an artifact of differences in firing rate.

The spike-count correlation might have become negative during saccades to the jointly preferred location because of uncontrolled trial-to-trial variation of a covariate for which the paired neurons had opposite selectivity. For example, if the firing rate of one neuron increased and the firing rate of the other neuron decreased with increasing saccadic amplitude, and if saccadic amplitude varied slightly from trial to trial, then a negative spike-count correlation would arise trivially from their opposed amplitude selectivity. To minimize any such artifact, we based all of the preceding analyses on residuals remaining after we had factored out the dependence of each neuron’s firing rate on covariates including the sequential position of the trial in the run, the duration of the delay period, the saccadic reaction time, the saccadic peak velocity, the direction and amplitude of the saccade, the initial angle of gaze and the final angle of gaze. The factoring procedure assumed, however, a linear dependence of firing rate on each covariate. In the event of nonlinear dependence, some influence of the covariate might have persisted in the residuals and have given rise to an artifactually negative spike-count correlation. To rule out this possibility, we assessed the impact of two manipulations on the tendency for the spike-count correlation to assume a negative value during saccades to the location preferred by neurons with strong and matching spatial preferences (spatial match index ≥ 0.4). First, we omitted the initial step of factoring out the dependence of firing rate on the covariates. If the negative r_sc_ were a covariate artifact, we would expect this manipulation to increase the negative magnitude of the median r_sc_. Contrary to this expectation, the magnitude was greater when the influences of all covariates had been factored out (“all” in Figure 6) than when the influence of no covariate had been factored out (“none” in Figure 6) or when any single covariate had been spared from the factoring process (intermediate bars in Figure 6). The fact that the magnitude was greater with the influences of all covariates factored out than under any other condition presumably is due to factoring having reduced the total variance and so increased the fraction of variance explained by genuine cross-neuronal covariation. Second, we reinstated the initial step of factoring out the dependence of firing rate on the covariates but we split neuron pairs into two categories based on whether their initial dependence on a covariate was of the same or opposite sign. Insofar as the measured r_sc_ was a covariate artifact, we would expect r_sc_ to be negative among opposite-sign pairs but positive among same-sign pairs. Contrary to this expectation, the measured median r_sc_ was negative among pairs in both categories regardless of whether the opposite-sign same-sign categorization was based on any single covariate or on the ten-dimensional vector representing combined dependence on all covariates.

**Figure 6.**
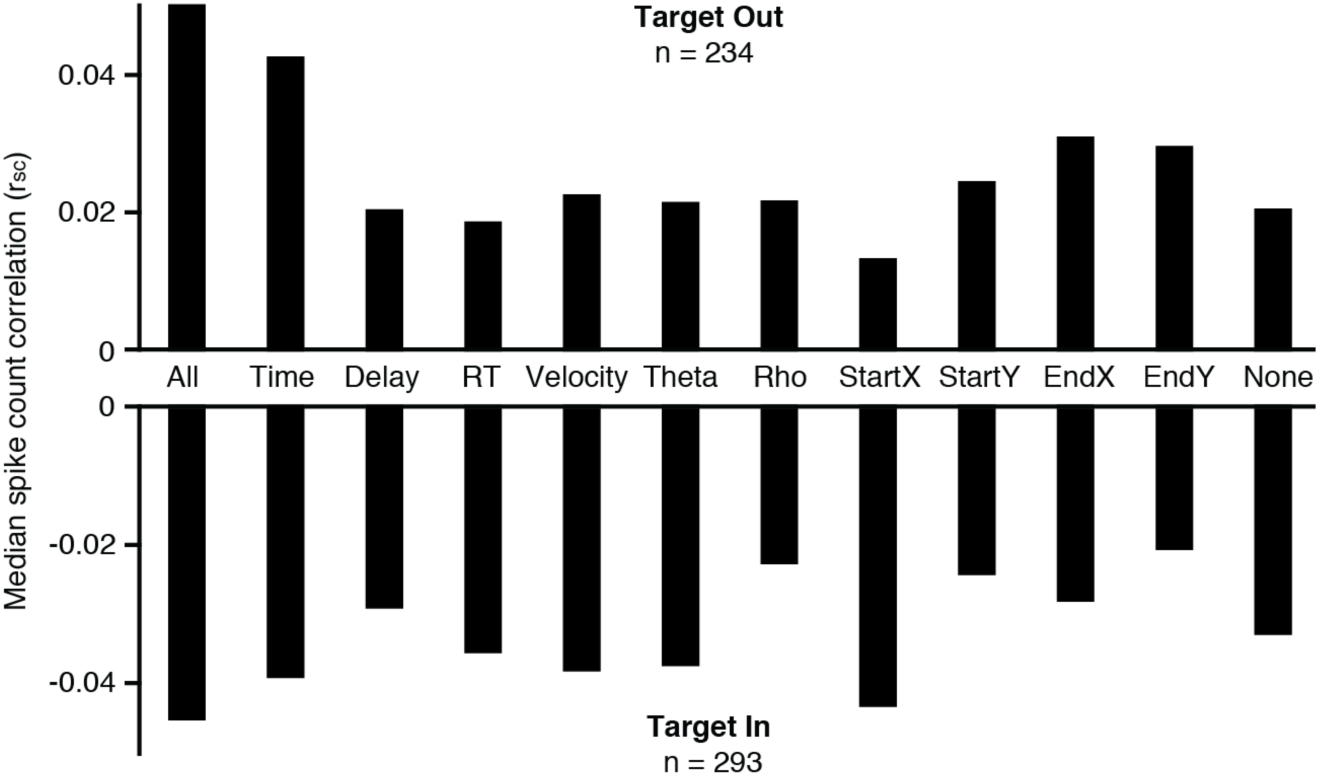
The push-pull effect did not arise from dependence of neuronal activity on behavioral or contextual covariates. Bar height indicates the median spike-count correlation in an analysis conducted after factoring out cross-trial variance in each neuron’s firing rate explained by the indicated covariate. The leftmost bars (“all”) represent results obtained after factoring out neuronal activity dependent on all of covariates. The rightmost bars (“none”) represent results obtained without factoring out neuronal activity dependent on any covariate. The intermediate bars represent results obtained after factoring out neuronal activity dependent on individual covariates. Time: the sequential number of the trial within the data collection session. Delay: the duration of the delay period as selected randomly on each trial. RT: Saccadic reaction time. Velocity: peak velocity of the eye. Theta and Rho: polar coordinates of the saccadic vector. Start X and Start Y: pre-saccadic gaze direction in Cartesian coordinates. End X and End Y: post-saccadic gaze direction in Cartesian coordinates. The analysis was based on neuron pairs above spatial-match threshold (d’•d’ > 0.4) which also met the conditions for computing a spike-count correlation in a 200 ms window centered on saccade initiation under the specified target condition. Each n indicates the number of such pairs.

The above analyses focused on neuron pairs preferring the same target location. To determine whether comparable phenomena occurred for neuron pairs with mismatched spatial preferences, we carried out a set of analyses identical to those described above but focused on neuron pairs with negative spatial match indices. We found that the median spike-count correlation at the time of the saccade was statistically indistinguishable from zero (Figure 7, *A*) and that there was no phasic change in the spike-count correlation around the time of the saccade (Figure 7, *B*).

**Figure 7.**
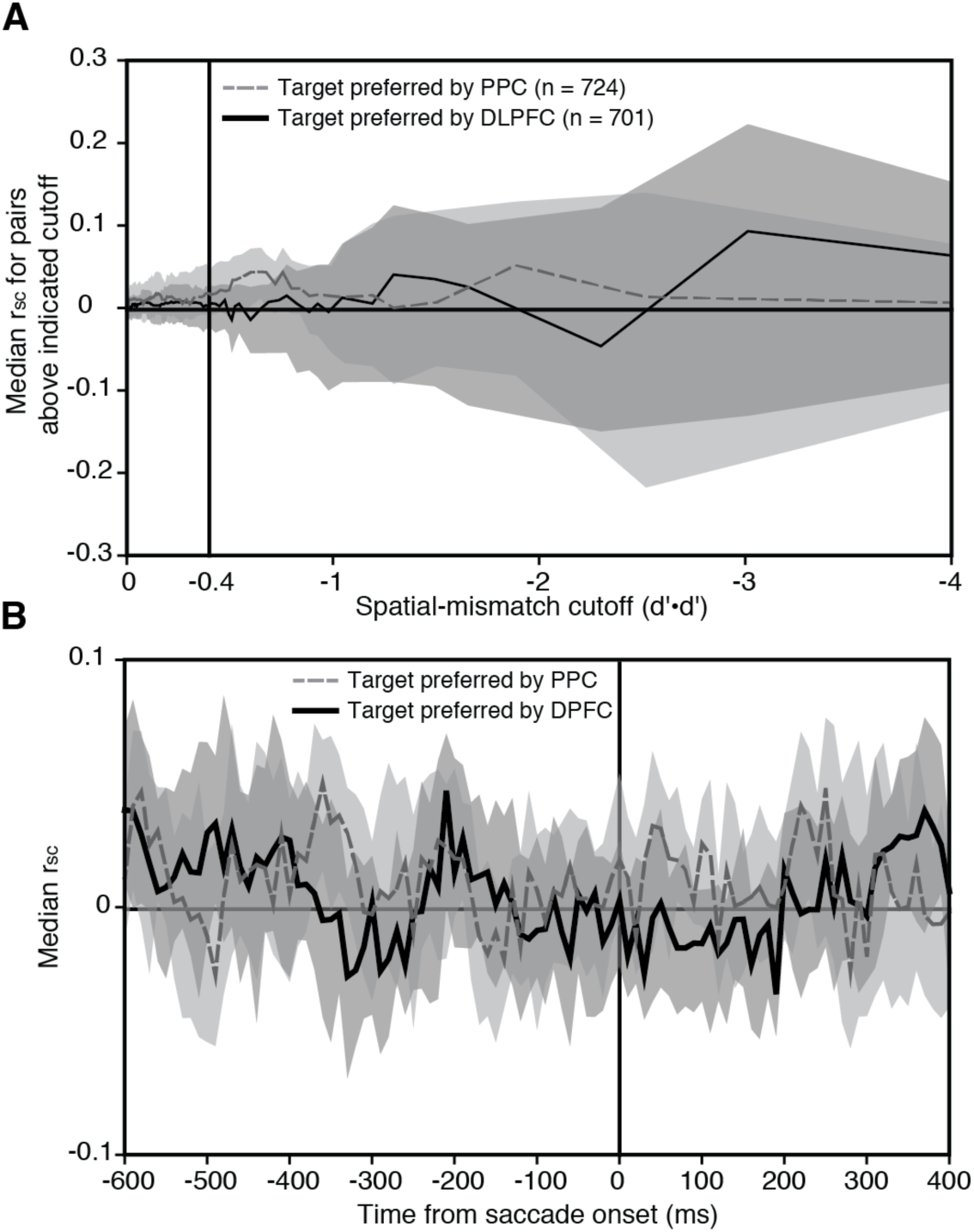
There was no push-pull effect in neuron pairs with mismatched location preferences. ***A***, The median spike-count correlation of DLPFC-PPC neuron pairs was statistically indistinguishable from zero. ***B***, This was true throughout the period preceding and following the saccade. Conventions as in Figure 4 with the sole exception that the horizontal axis in ***A*** contains values in the negative range, with the strongest degree of spatial mismatch represented farthest to the right.

Previous studies of neuron pairs within the same area have established that the spike-count correlation is positive on average regardless of trial epoch in both DLPFC and PPC (Constantinidis and Goldman-Rakic, 2002; Cohen et al., 2010; Qi and Constantinidis, 2012; Leavitt et al., 2013; Katsuki et al., 2014b; Leavitt et al., 2017b, a). To determine whether this was true in our study, we carried out parallel analyses on data from 1281 DLPFC-DLPFC pairs involving 454 neurons and 1252 PPC-PPC pairs involving 414 neurons. The analyses necessarily were restricted to neurons with matched spatial preferences because neurons recorded on the same linear microelectrode array nearly always had congruent spatial selectivity. In both DLPFC and PPC, the within-area median spike-count correlation was strongly positive throughout the analysis period. In DLPFC, the magnitude of the correlation appeared not to vary as a function of target location (Figure 8, *A*) or time relative to saccade onset (Figure 8, *B*). In PPC, it was higher under the target-out than under the target-in condition (Figure 8, *C*) specifically during the period immediately before saccade onset (Figure 8, *D*).

**Figure 8.**
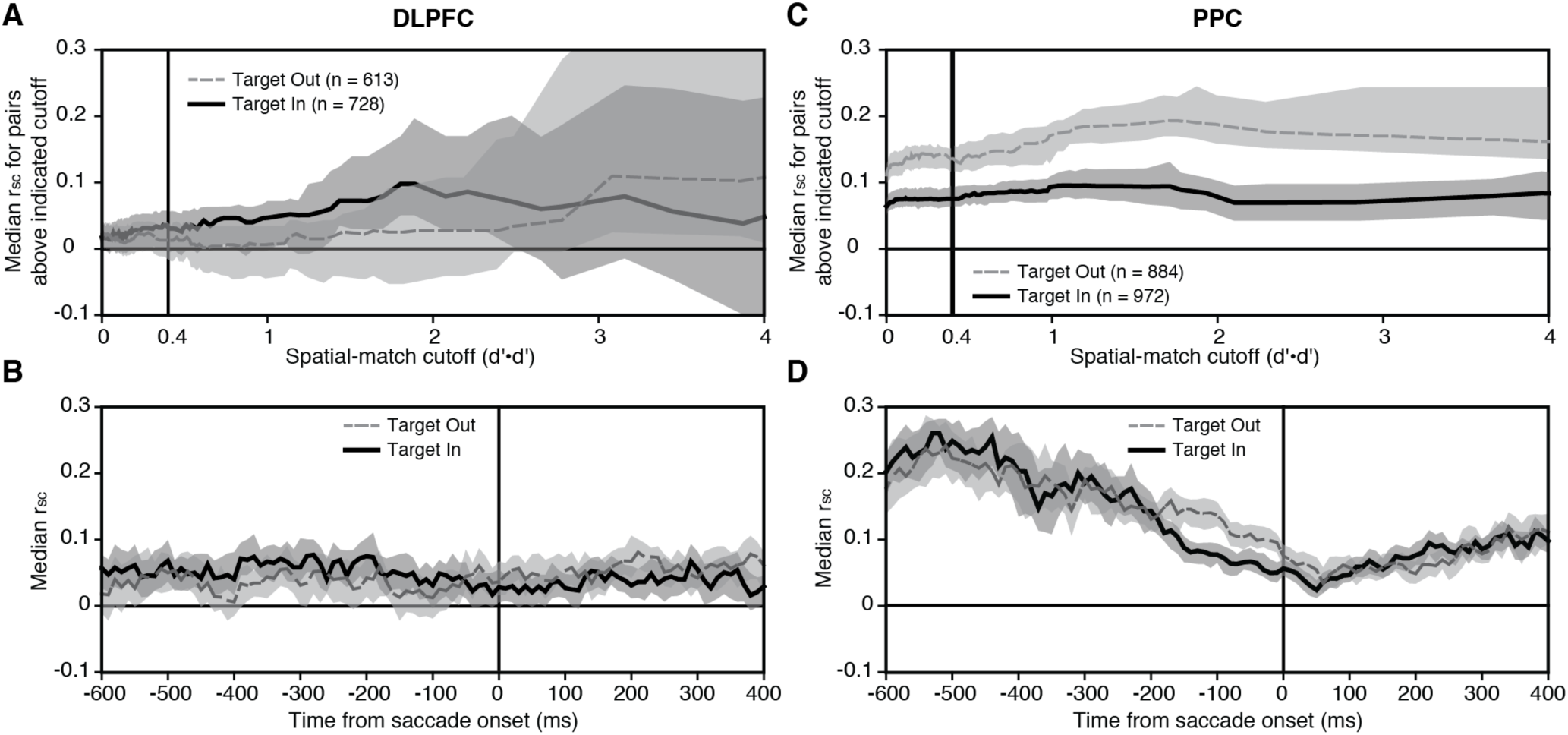
There was no push-pull effect between neurons in the same area. ***A***, For DLPFC-DLPFC neuron pairs with matched spatial selectivity, the median spike-count correlation was slightly positive under both the target-in and the target-out conditions. ***B***, This was true throughout the period preceding and following the saccade. ***C***, For PPC-PPC neuron pairs with matched spatial selectivity, the perisaccadic spike-count correlation was reduced under the target-in condition as compared to the target-out condition. Analysis based on a 200 ms window centered 100 ms before saccade initiation. ***D***, This effect was confined to a period of around 200 ms preceding and accompanying the saccade. All conventions as in Figure 4.

## Discussion

The key finding of this study is that under certain well defined conditions the spike-count correlation between prefrontal and parietal neurons shifts from positive to negative. The necessary conditions are that the neurons have matching spatial preferences and that a saccade be directed into the joint response field. The excursion into negativity is brief, being confined to the time of saccade execution. This is the first instance in which neurons in different cortical areas have been demonstrated to exhibit predominantly negative spike-count correlations. The existence of a negative correlation implies that prefrontal and parietal neurons contributing to the execution of a saccade become subject to some competitive process around the time of the saccade. We cannot be certain of the functional significance of this phenomenon. We note, however, that it can be accommodated within the framework of optimal feedback control theory (Todorov and Jordan, 2002; Pruszynski and Scott, 2012). Optimal feedback control is possible in any system containing effectors with redundant actions. The key principle of optimal feedback is that noise at the level of the effectors should be controlled only to the degree that it impairs achievement of a defined goal. Maintaining constant water flow under control of hot and cold taps is a simple example (Pruszynski and Scott, 2012). An optimal controller ensures that the sum of the two settings is constant without regard to the individual settings. As an incidental consequence of this arrangement, if the individual settings vary over time, they do so in a negatively correlated pattern. The negative spike-count correlation between prefrontal and parietal neurons might, by analogy, emerge in a system obeying the constraint that saccade bursts converging on the superior colliculus from multiple cortical areas sum to a constant value. This constraint is in harmony with the observation that each saccade, regardless of its amplitude or direction, is associated with activity in a collicular burst zone of stereotyped extent and magnitude (Munoz and Wurtz, 1995).

In numerous previous studies of neuron pairs in the same cortical area, the spike-count correlation has always been observed to be positive on average (Cohen and Kohn, 2011). The strength of the positive correlation is, however, lower for pairs that are far apart (Constantinidis and Goldman-Rakic, 2002; Smith and Kohn, 2008; Cohen et al., 2010; Leavitt et al., 2013; Smith and Sommer, 2013; Ecker et al., 2014; Katsuki et al., 2014b) or have discordant patterns of selectivity (Zohary et al., 1994; Bair et al., 2001; Constantinidis and Goldman-Rakic, 2002; Cohen and Newsome, 2008; Smith and Kohn, 2008; Cohen et al., 2010; Gu et al., 2011; Hansen et al., 2012; Qi and Constantinidis, 2012; Leavitt et al., 2013; Smith and Sommer, 2013; Ecker et al., 2014; Ruff and Cohen, 2014b; Markowitz et al., 2015; Chelaru and Dragoi, 2016; Leavitt et al., 2017b, a) and may vary as a function of wakefulness (Ecker et al., 2014), effort (Ruff and Cohen, 2014a), attention (Cohen and Maunsell, 2009; Mitchell et al., 2009; Herrero et al., 2013; Luo and Maunsell, 2015; Ni et al., 2018), learning (Cohen and Newsome, 2008; Cohen et al., 2010; Gu et al., 2011; Qi and Constantinidis, 2012; Ruff and Cohen, 2014b; Markowitz et al., 2015; Ni et al., 2018) and task set (Cohen and Newsome, 2008; Cohen et al., 2010; Ruff and Cohen, 2014b, a). Although centered in the positive range, the distribution of spike-count correlations typically extends into the negative range. Significant negative correlations, observed to occur more frequently than expected by chance in studies of V1 (Hansen et al., 2012; Chelaru and Dragoi, 2016), MSTd (Gu et al., 2011), FEF (Cohen et al., 2010), area 8a (Leavitt et al., 2013) and PFC (Markowitz et al., 2015), tend to occur under conditions otherwise associated with low positive correlations, for instance between neuron pairs that are far apart (Cohen et al., 2010; Leavitt et al., 2013) or that have opposed patterns of spatial selectivity (Cohen et al., 2010; Hansen et al., 2012; Leavitt et al., 2013; Chelaru and Dragoi, 2016). The push-pull phenomenon we have described is clearly different from within-area interactions insofar as it involves a competitive interaction between neurons with matched spatial selectivity.

Few previous studies have characterized spike-count correlations between neurons in different cortical areas. Two recent cases concerned paired recording in areas V1 and MT (Ruff and Cohen, 2016a, b, 2017) and in areas V1 and FEF (Pooresmaeili et al., 2014). In both cases, the spike-count correlation of neuron-pairs with overlapping response fields was positive on average and correlation strength increased with attention to an image located in the zone of overlap. This outcome may reflect a principle whereby attention enhances functional connectivity between neurons representing image content at the attended location (Ruff and Cohen, 2016b). The current results cannot be accommodated in this framework because under conditions requiring attention to the location of the target, the spike-count correlation of neurons representing that location undergoes an excursion into the negative range indicative of inverted functional connectivity. Our findings instead suggest a fundamental distinction between area-area interactions involving the visual system, where neurons representing the same stimulus engage in cooperative interaction once it has been selected for attention, and in the executive control system, where neurons representing the same action engage in competitive interaction once it has been selected for execution.

## Acknowledgements

We thank Douglas Ruff for his comments on data analysis and presentation. We thank Karen McCracken for technical assistance. Support from NIH RO1 EY024912 and P50 MH103204. Technical support from NIH P30 EY008098.

The authors declare no competing financial interests.

